# Coevolution of larval signalling and worker response can trigger developmental caste determination in social insects

**DOI:** 10.1101/2023.09.27.559532

**Authors:** Juan J. Lagos-Oviedo, Ido Pen, Jan J. Kreider

## Abstract

Eusocial insects belong to distinct queen and worker castes, which, in turn, can be divided into several morphologically specialised castes of workers. Caste determination typically occurs by differential nutrition of developing larvae. We present a model for the coevolution of larval signalling and worker task allocation – both modelled by flexible smooth reaction norms – to investigate the evolution of caste determination mechanisms and worker polymorphism. In our model, larvae evolve to signal their nutritional state to workers. The workers evolve to allocate time to foraging for resources versus feeding the brood, conditional on the larval signals and their body size. Worker polymorphism evolves under accelerating foraging returns of increasing body size, which causes selection to favour large foraging and small nursing workers. Worker castes emerge because larvae evolve to amplify their signals after obtaining some food, which causes them to receive more food, while the other larvae remain unfed. This leads to symmetry-breaking among the larvae who either are well-nourished or malnourished; thus, emerging as small or large workers. Our model demonstrates the evolution of nutrition-dependent caste determination and worker polymorphism by a self-reinforcement mechanism that evolves from the interplay of larval signalling and worker response to the signals.

## 1. Introduction

The emergence and evolution of polymorphism is a central topic in evolutionary biology [1,2]. Polymorphism is found both at the genetic [3] and organismic level [4]. An extreme case of polymorphism is prevalent in eusocial insects in which workers within a colony can vary highly in body size [5–8]. Bumblebee workers of the same colony, for instance, exhibit up to tenfold difference in body weight [9]; and around 30% of ant species have workers with large variation in body size [10], who may belong to distinct size classes and thus form distinct worker castes [5–7]. Worker castes are typically specialised in particular tasks within the colony, such as colony defence [11], foraging and food transport [12,13], brood care [14,15], or food storage [16,17]. Understanding the evolution of worker polymorphism and division of labour is a pivotal step towards comprehending the social organization of eusocial organisms.

Several candidate factors that might facilitate the evolution of worker polymorphism have been proposed [18]. The developmental hypothesis for the evolution of worker polymorphism suggests that worker polymorphism is more likely to evolve if queen-worker developmental pathways diverge relatively early during larval development because such an early divergence would leave a longer time period for a novel developmental divergence between distinct worker castes [19]. Furthermore, worker polymorphism might evolve more readily in species with large colonies because in small colonies investment in a few large specialised workers might be too risky compared to investment into more small workers [5,20]. Lastly, higher queen mating frequencies could facilitate the evolution of worker polymorphism by increasing genetic diversity among workers, thus leading to workers of more variable body sizes [18].

Apart from these potential facilitatory factors for the evolution of worker polymorphism, division of labour models typically predict that task specialisation and division of labour evolve if there are efficiency benefits of specialisation [21–23]. Morphological differences can cause some individuals to be more efficient in some tasks than others [24]; for instance, in various species of ants larger workers are more efficient in carrying food [13,25–27], defending the nest [11], or dealing with heat [28,29].

In social insects, the body size of adult individuals is typically strongly influenced by food quality and quantity obtained during larval development [6,30–35]. This is most extreme in species that have morphologically distinct, specialised castes, where caste differentiation is often achieved by differential nutrition during larval development [6,30,31,34–37]. In an established colony, workers control the amount of food that particular larvae obtain [38], and larvae can communicate their nutritional requirements to the workers by producing larval signals [39,40], mediated pheromonally [41,42] or behaviourally [43–45]. The worker body size distribution within a colony might thus be driven by a complex interplay between larval signalling and worker response to the larval signals. However, the outcome of this complex interplay is difficult to understand intuitively; hence, the need for a formal model of the coevolution of larval signalling and worker behaviour.

Here, we present an individual-based simulation model for the coevolution of larval signalling behaviour and worker task choice. In the model, larvae can evolve to signal their hunger state to the workers. The amount of resources a larva obtains determines its body size when it develops into a worker. Workers, in turn, can evolve to allocate time to foraging for resources or nursing the brood, depending on their body size and their perception of the larval signals. We model these condition-dependent behaviours with highly flexible reaction norms to allow for the evolution of non-obvious, non-linear strategies. We evaluate the effect of the duration of development, queen mating frequency, and costs for larval signal production on the evolution of worker polymorphism.

## 2. Methods

### (a) General model setup

We developed an evolutionary individual-based simulation model representing a population of a fixed number of *N* social insect colonies (parameter values in Table 1). In the model, colonies are independently founded by queens who produce larvae that develop into workers or queens. Larvae can evolve to signal their nutrition level to workers. Workers evolve to allocate time to foraging or nursing larvae depending on their body size and the larval signals (Fig. 1).

**Table 1.** Parameter values used in the simulations, unless stated otherwise.

| Variable | Value | Meaning |
| --- | --- | --- |
| $N$ | 1000 | Number of colonies |
| $X_{\max}$ | 5.0 | Upper body size limit |
| $C$ | 0.0 | Signalling cost |
| $m$ | 0.05 | Mutation rate |
| $\mu$ | 0.0 | Mutation mean |
| $\sigma_1$ | 0.01 | Mutational standard deviation for reaction norm parameters |
| $\sigma_2$ | 0.2 | Mutational standard deviation for $\alpha$ (eq. 3) |
| $T_{\text{season}}$ | 50 | Number of time steps per season |
| $T_{\text{repro}}$ | 5 | Number time steps of reproductive phase |
| $T_{\text{dev}}$ | 4 | Developmental time |
| $n$ | 1 | Queen mating frequency |
| $r$ | 0.5 | Proportion of resources invested in reproduction |
| $\mu_{\text{feed}}$ | 0.5 | Mean resources transferred while feeding |
| $\sigma_{\text{feed}}$ | 0.2 | Standard deviation of resources transferred while feeding |
| $\mu_{\text{egg}}$ | 0.5 | Mean resources transferred through egg |
| $\sigma_{\text{egg}}$ | 0.2 | Standard deviation resources transferred through egg |
| $a$ | 0.2 | Scaling parameter for foraging function |
| $b$ | 1.5 | Scaling parameter for foraging function |
| $f$ | 12.0 | Scaling parameter for larval mortality function |
| $g$ | -8.0 | Scaling parameter for larval mortality function |

**Figure 1.**
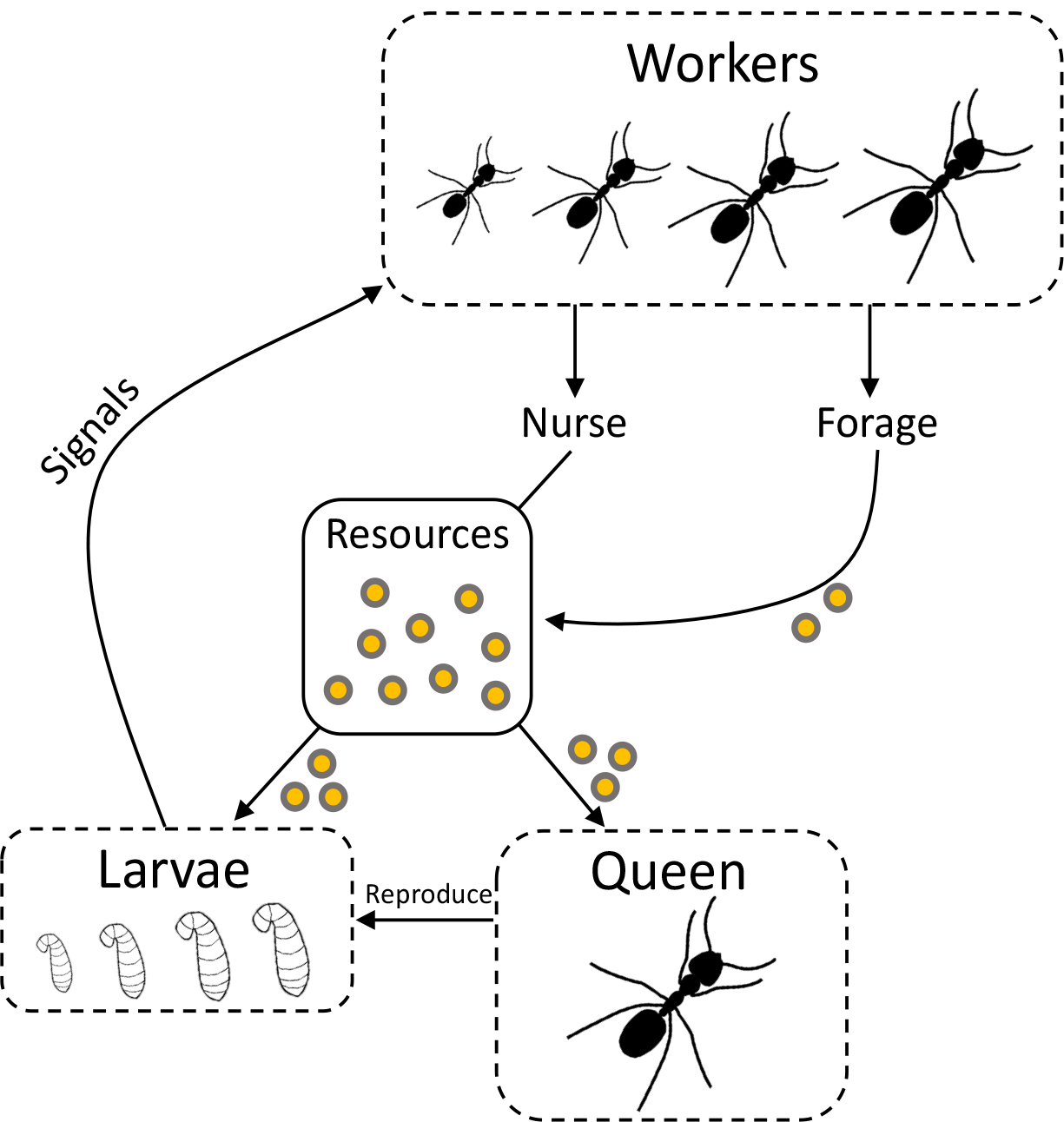
Task allocation and resource flow within a colony. Workers vary in their body size, and allocate time to foraging or nursing. Workers evolve a condition-dependent task choice which is a function of their body size and the level of larval signals. If workers forage, they obtain resources (yellow circles) that are stored in the resource stock of the colony (centre). If workers nurse, they take some of the resources from the resource stock to feed the larvae. The body size of workers developing from the larvae is proportional to the amount of resources they obtained. The queen uses some of the resources from the resource stock to reproduce, thus increasing the number of larvae. Larvae can evolve to signal their nutrition level to the workers.

### (b) Larval signalling

Larvae have an evolving smooth reaction norm modelled by a natural cubic spline that determines the larval signalling strength by a larva as a function of its nutrition level. Natural cubic splines are functions consisting of connected cubic polynomials, which allows the functions to take highly flexible shapes (though both ends are linear; for details, see Supplement). After having signalled, the larva’s nutrition level decreases by the product of the signalling strength and the signalling cost *C*. At the start of the simulation, larvae produce no signals, regardless of their nutrition level, but they can subsequently evolve to produce a nutrition-dependent signal.

### (c) Development

After *T*_dev_ time steps, a larva may develop into a worker. The nutrition level of a larva determines its body size when it develops into a worker; thus, the amount of food obtained as a larva scales linearly with worker body size. The body size of workers *X* has an upper limit of *X*_max_. The probability that the larva successfully develops increases with the amount of food that it obtained. The probability that the larva does not develop and dies is modelled by a logistic function

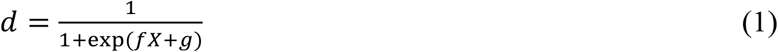

where *f* and *g* are parameters that affect the function’s steepness and location.

### (d) Task choice

The propensity of a worker to forage is determined by a reaction norm modelled by a natural cubic spline surface (for details, see Supplement). This reaction norm is a function of the sum of larval signals produced by the entire colony and the worker’s body size. If workers do not forage, they nurse the larvae. At the start of the simulation, the reaction norm is flat, resulting in a foraging probability of 0.5, independently of larval signals or the worker’s body size. Subsequently, the reaction norm can evolve, so that the foraging probability can come to depend on larval signals and/or worker body size.

### (e) Genetics

Individuals have a haplodiploid sex determination system; diploid females develop from fertilized eggs and haploid males from unfertilized eggs. Female individuals carry two sets of 31 genes, five of which determine the shape of the reaction norm for larval signalling, 25 of which determine the shape of the reaction norm for the propensity of a worker to forage (for details, see Supplement), and one of which determines the worker’s ability to selectively feed larvae with stronger signals (see below). The genes are assumed to be unlinked; hence they can recombine freely. Workers randomly express either the maternal or paternal copy of the homologous genes on a per-gene basis. Gene values mutate with a per-locus mutation rate of *m* each meiotic event. Fig. S1 shows how different parameter values for *m* affect the results. Mutational effect sizes for the evolving reaction norm and the ability of workers to selectively feed larvae (see below) parameters are drawn from normal distributions.

### (f) Life cycle

Colonies have a seasonal semelparous life cycle with *T*_season_ time steps per generation. Each generation is composed of a founding phase, a growth phase, and a reproductive phase. During the founding phase, a single founding queen establishes a colony. The founding queen forages, reproduces, and feeds the larvae. We assume that queens obtain one resource unit during foraging and subsequently feed one larva per time step. The founding phase ends as soon as the first worker is present. The colony then enters the growth phase, during which the queen continues laying eggs but stops foraging and feeding larvae, while the workers forage or nurse. During the last phase of the season, the colony enters the reproductive mode, which takes *T*_repro_time steps. The larvae now develop into queens and males instead of workers. Before entering hibernation, queens mate with *N* males, randomly selected from the population as a whole, and store their sperm for use in the next season. We assume that offspring are produced with an even sex ratio in the reproductive phase. Males are only produced in the reproductive phase. Before the subsequent generation emerges, the population size is maintained by randomly sampling *N* of the mated hibernating queens.

### (g) Foraging

Foraging workers acquire resources and carry them to the colony, where they are stored in the colony resource stock of variable size *R*. The foraging return of a worker of body size *X* is given by the power function

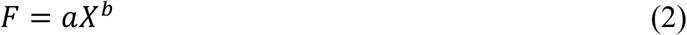

where *a* is a parameter that scales the function and *b* is a parameter that determines the function shape. If *b* < 1, then foraging has diminishing returns with increasing body size. If *b* = 1, then foraging returns increase linearly with increasing body size. If *b* > 1, then foraging returns accelerate with increasing body size.

### (h) Nursing

Founding queens and nursing workers feed resources to the larvae from the colonies’ resource stock. Feeding increases a larva’s nutrition level. The probability of larva *l* to be fed is given by the *softmax* function

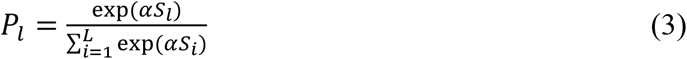

where *S*_*l*_ is the signalling strength of larva *l* and *S*_*i*_ the signalling strength of larva *i*, which ranges from 1 to the number of larva *L*. The evolving parameter *α* ≥ 0 describes the nurses’ (or founding queen’s) ability to discriminate and selectively feed larvae. If *α* = 0, then larvae are fed randomly and thus independently of their relative signalling strengths. The larger *α*, the greater the share of food obtained by larvae with stronger signals. Fig. S2 illustrates the feeding probabilities that different values of *α* results in. Nurses and founding queens transfer an amount of resources to the larvae, sampled from a zero-truncated normal distribution with mean μ_feed_ and standard deviation σ_feed_. We assume that feeding in the reproductive phase is always random with respect to signalling strength of the larvae.

### (i) Reproduction

Every time step, the queen invests a proportion *r* of the *R* resources stored in the colony into reproduction. Consequently, the reproduction of the queen scales linearly with the amount of resources in the colony, imposing strong selection on worker task allocation to foraging vs. nursing the brood. We assume that egg size varies randomly to a small extent relative to the mean egg size. Thus, larvae obtain resources from the egg sampled from a zero-truncated normal distribution with mean μ_egg_ and standard deviation σ_egg_. The queen lays eggs until the resources used for reproduction, *rR*, are depleted. Fig. S3 shows how different parameter values for *r* affect the results.

### (j) Model analysis

We implemented the model in C++, compiled it with g++ 11.2.0., and analysed and visualised model results in R 4.2.2 [46] using the packages *tidyverse* [47], *MetBrewer* [48], *cowplot* [49], and *ggpubr* [50]. We simulated the model for 5000 generations. For each replicate simulation, we assessed the proportion of colonies with a multimodal body size distribution by applying Hartigan’s dip test [51] to the body size distribution within a colony using the R-package *diptest* [52]. We rejected the assumption of unimodality at a significance level of 0.1. In the figures, we report the percentage of colonies with polymorphic workers of 50 randomly sampled colonies (due to computational constraints), i.e., the proportion of colonies for which the assumption of a unimodal body size distribution could be rejected.

## 3. Results

### (a) Worker caste evolution

Colonies evolve to have polymorphic worker castes only when foraging returns accelerate with increasing body size but not under diminishing or linear returns (Fig. S4). To elucidate the mechanisms by which polymorphism among workers emerge, we compare simulations with diminishing foraging returns (*b* = 0.5) to simulations with accelerating returns (*b* = 1.5).

Under diminishing returns, the larvae evolve a monotonically increasing signalling reaction norm and the worker body size distribution evolves to be unimodal (Fig. 2a + b). Workers evolve a relatively weak preference to feed (*α* = 1.43 mean ± 0.02 se) more strongly signalling larvae. Workers evolve to allocate relatively more time to nursing when larval signalling levels are high, i.e., when there are many larvae. However, at low and intermediate levels of larval signalling, small workers evolve to prefer nursing whereas larger workers are more likely to forage (Fig. 2c). Consequently, some level of size-based task specialisation evolves.

**Figure 2.**
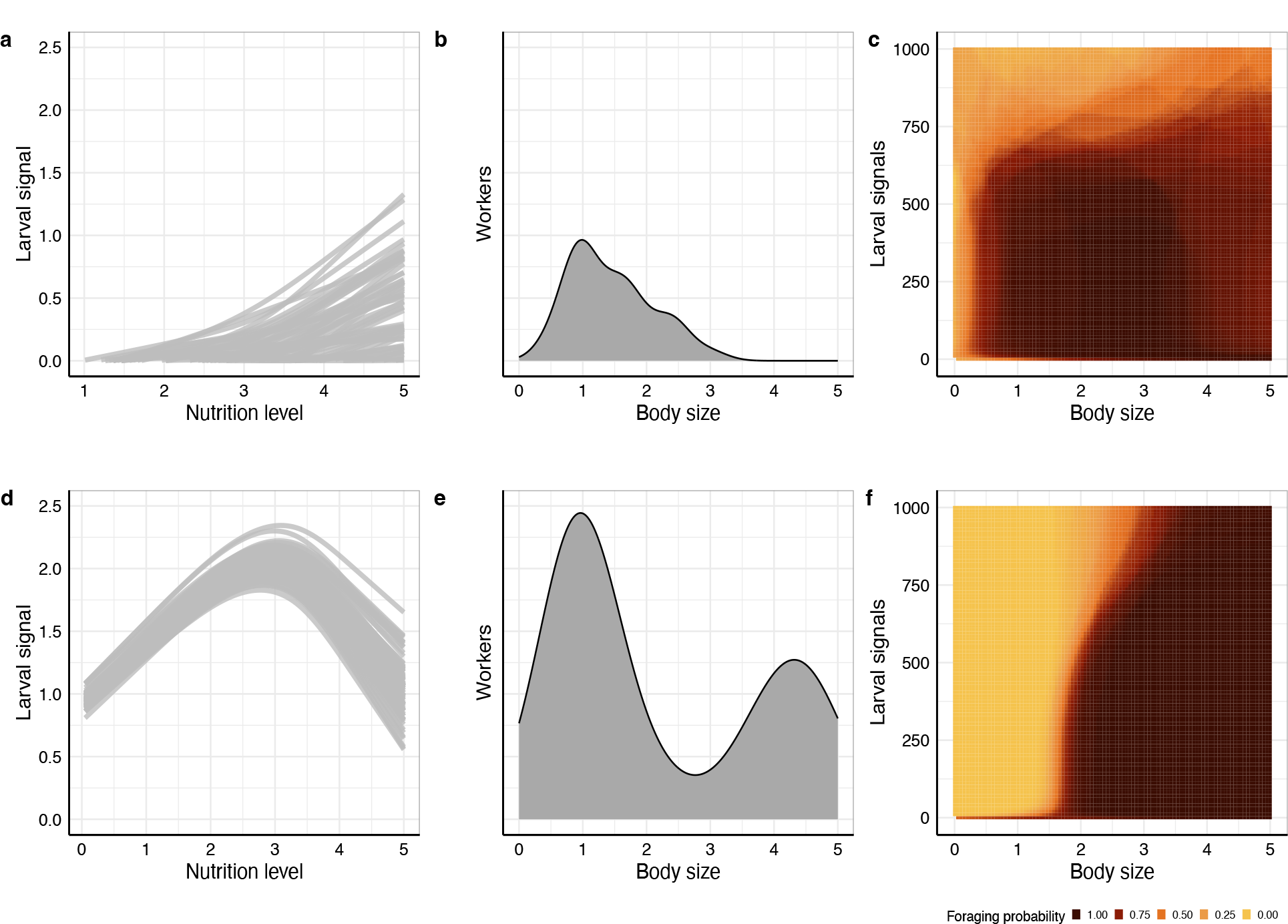
Evolved larval signalling reaction norms, body size distributions, and worker task allocation reaction norms. (a - c) Simulations with diminishing foraging returns of increasing body size (*b* = 0.5). (d - f) Simulations with accelerating foraging returns of increasing body size (*b* = 1.5). (a, d) Evolved larval signalling as a function of larval nutrition level for 100 random larvae. Each grey line represents the evolved reaction norm of a larva. (b, e) Worker body size distribution density plot for a random colony. (c, f) Evolved worker foraging probability as a function of the sum of larval signals and worker body size. Colours show the mean of the evolved reaction norms of 100 random workers.

In contrast, under accelerating returns, distinct body size classes of smaller and larger workers that specialise on nursing the brood and foraging, respectively, emerge, while workers of intermediate sizes are apparently disfavoured. Larvae evolve hump-shaped reaction norms with a maximum at intermediate nutrition levels (Fig. 2d). As a result, larvae that have been fed once are more likely to be fed again until their nutrition level exceeds some intermediate level. Subsequently, their signalling level decreases up to a point where larvae with considerably lower nutrition levels signal more and are more likely to receive food, thus preventing resources from being wasted on well-nourished larvae that cannot increase their size further. As a consequence, larvae either have a low nutrition level, if they – by chance – did not receive food initially, or a high nutrition level, if they did receive some food, which increases their probability to receive even more food. Small initial differences in the larvae’s nutrition level originating from differences in egg size can facilitate the differentiation of nutrition levels of different larvae. However, they are not necessary for symmetry-breaking among the larvae because worker polymorphism also evolved in the absence of any differences in egg sizes (Fig. S6). The reaction norm shape in Fig. 2d represents a self-reinforcement mechanism which yields two distinct groups of larvae: those with low and those with high nutrition levels. In addition, workers evolve a relatively strong preference (*α* = 5.04 mean ± 0.01 se) to feed “louder” larvae. Thus, the interplay of the larval signalling reaction norm and the worker’s feeding preference results in a bimodal body size distribution of workers (Fig. 2e). Moreover, workers evolve a reaction norm such that specialisation by body size occurs, with large workers foraging and small workers nursing; thus, division of labour evolves (Fig. 2f).

### (b) Developmental time

It has been hypothesised that worker polymorphism is more likely to evolve if a longer time period is available for a developmental divergence between distinct worker castes [19]. To investigate this, we varied the duration of development in our model. If developmental times are shorter, then fewer colonies evolve to have polymorphic workers (Fig. 3). Since the self-reinforcement mechanism that leads to the emergence of polymorphic workers in our model is based on multiple feeding events of the same individual larva, longer developmental periods can indeed favour the emergence of polymorphic workers, because longer developmental periods increase the number of times a larva can receive food. However, the facilitatory effect of developmental length is only quantitative, because at least some colonies with polymorphic workers always occur, regardless of the duration of development.

**Figure 3.**
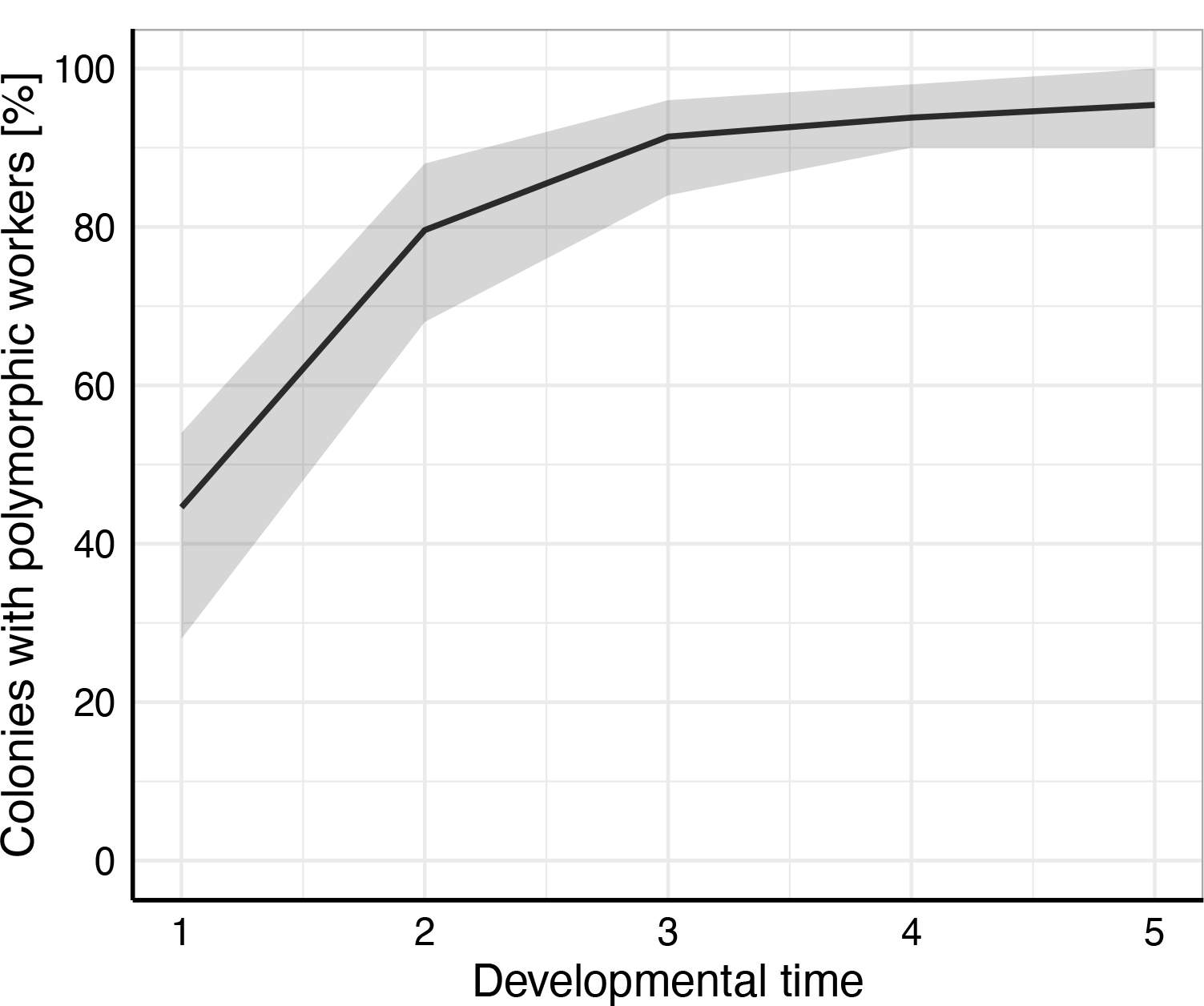
The effect of the length of the developmental period on the evolution of worker polymorphism. We simulated a range of 1 to 5 in steps 1 of developmental time (*T*_dev_), each with 10 replicate simulations. Black lines show the mean percentage of colonies with polymorphic workers and grey areas the range across replicates. We assume accelerating foraging returns of increasing body size (*b* = 1.5).

### (c) Queen polyandry

It has been suggested that high levels of queen polyandry could facilitate the evolution of worker polymorphism due to increased within-colony genetic variation [18]. We therefore investigated the effect of queen mating frequency on the evolution of worker polymorphism. In our model, queen mating frequency had barely any effect on the evolution of worker polymorphism (Fig. S5).

### (d) Larval signalling cost

The production of larval signals is often assumed to be associated with some energetic cost [53,54]. As signal production becomes more costly, the proportion of colonies with polymorphic workers decreases until worker polymorphism does not evolve at all (Fig. 4). This is because signalling does not evolve if it is too costly. Consequently, the self-reinforcement mechanism for the emergence of worker polymorphism does not evolve either.

**Figure 4.**
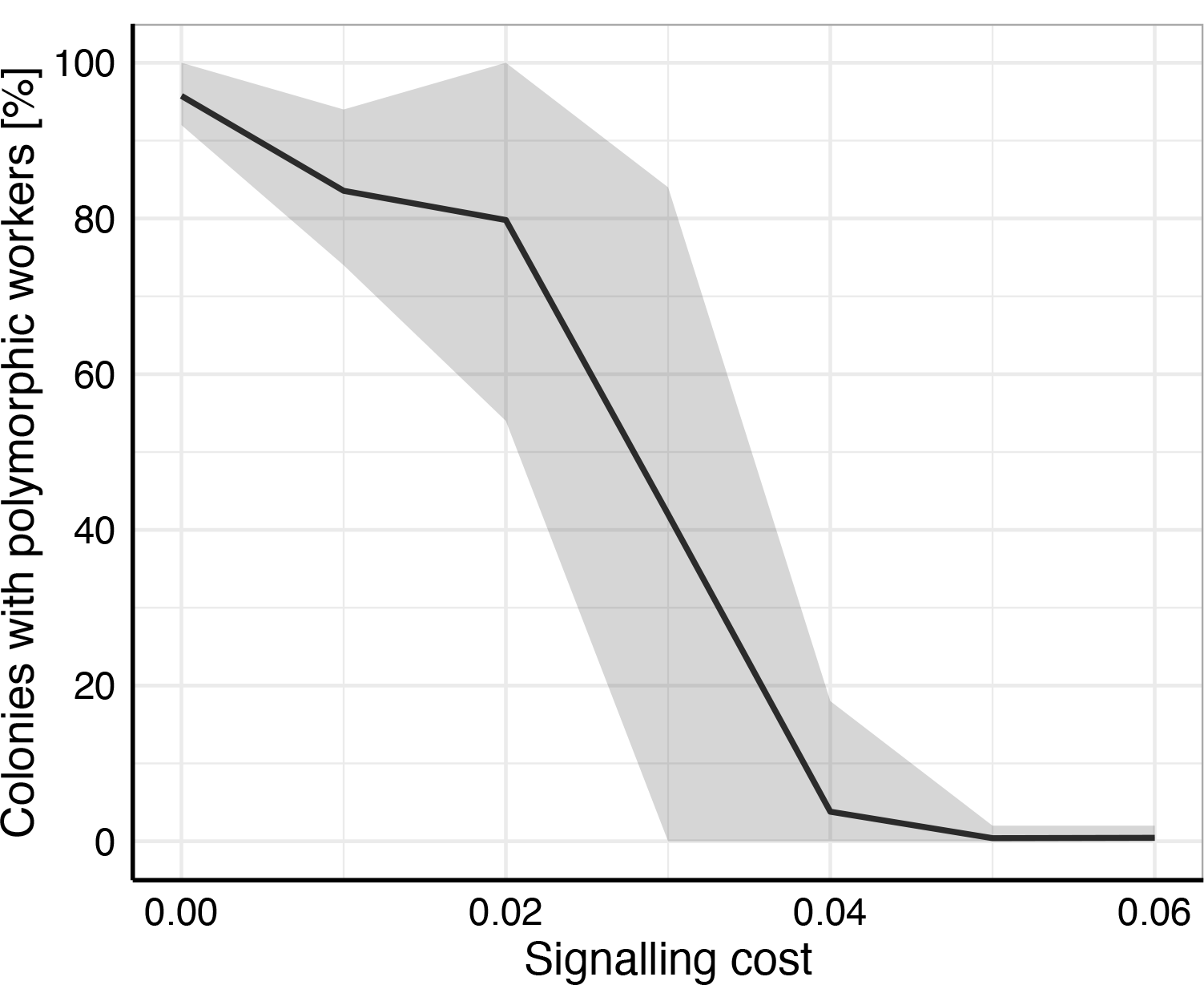
The effect of signalling costs on the evolution of worker polymorphism. We simulated a range of 0.00 to 0.06 in steps of 0.01 of signalling costs (*C*), each with 10 replicate simulations. For comparison, the amount of resources a larva obtains when being fed is μ_feed_ = 0.5 on average. Black lines show the mean percentage of colonies with polymorphic workers and grey areas the range across replicates. We assume accelerating foraging returns of increasing body size (*b* = 1.5).

## 4. Discussion

We here present a model for the coevolution of larval signalling and worker task allocation to investigate the evolution of worker polymorphism in eusocial insects. Our model demonstrates how mechanisms for symmetry-breaking among similar larvae and for worker caste determination can evolve, and thereby integrates evolutionary and mechanistic explanations [23].

### (a) Mechanisms of caste determination

The occurrence of morphologically and behaviourally distinct castes is one of the main characteristics of eusocial organisms [23,55–57]. In our model, worker castes of distinct body size emerge through an evolved self-reinforcement mechanism. This mechanism causes some larvae to develop into large workers and some larvae into small workers, even in the absence of any systematic differences between individual larvae, although random differences in egg size can facilitate the differentiation of nutrition levels between larvae. A role of self-reinforcement for the emergence of castes has also been demonstrated in a non-evolutionary division of labour model, where individuals with high nutrition levels are dominant over individuals with lower nutrition levels and thus obtain more resources, leading to the emergence of distinct groups of malnourished foragers and well-nourished nurses [58].

Across eusocial organisms, queen-worker and worker-worker caste differentiation typically occur by differential nutrition during larval development [6,30,31,34–37]. Although our model is not predisposed towards the production of workers of distinct size classes – there are no build-in size classes but body size is a continuous trait – we observe the emergence of polymorphic workers belonging to groups of distinct body sizes. This is because accelerating foraging returns with increasing body size cause disruptive selection to favour the production of small and large workers, while disfavouring workers of intermediate size. This, in turn, leads to the joint evolution of preferential feeding by the workers of larvae with stronger signals and a larval signalling strategy that has a maximum at intermediate nutrition levels. Thus, larvae signal more strongly after having obtained some initial amount of food, which makes them more likely to obtain food again, until their signalling level decreases at high nutrition levels. This decrease prevents resources from being wasted on larvae that are close to the maximum body size. The decrease seems realistic rather than being an artefact resulting from the limitation of body size because larval growth would also be limited, i.e., at some size, larvae would not grow further due to growth constraints, even when obtaining further food.

The evolved self-reinforcement mechanism that leads to polymorphic workers resembles a nutrition-dependent developmental switch for caste determination, where larvae develop into small workers after receiving only little food or into large workers when being fed a lot. Our model therefore demonstrates how nutrition-dependent caste determination – prevalent across eusocial organisms – can evolve mechanistically. In our model, also some workers of intermediate size emerge. This “bug” is actually a realistic feature, as workers of intermediate size also occur in polymorphic species [7] and queen-worker differentiation can sometimes lead to the production of intercaste individuals that share traits of both queens and workers [30].

### (b) Facilitation of worker polymorphism

Our model identifies a range of conditions under which the evolution of worker polymorphism is facilitated or hampered. Longer developmental periods promote the prevalence of worker polymorphism since longer developmental periods increase the number of times a larva can obtain food. This is in line with the developmental hypothesis for the evolution of worker polymorphism which suggest that a relatively early divergence of queen-worker developmental pathways could facilitate the evolution of worker polymorphism by leaving more time for a subsequent developmental divergence between different worker castes [19].

Queen mating frequency hardly has any effect on the prevalence of worker polymorphism under accelerating foraging returns of increasing body size. It has been suggested that high levels of queen polyandry could increase worker genetic diversity and consequently body size variation [18]. This is most likely to occur when there are direct genetic effects on body size [36,59,60], which are absent in our model.

Lastly, our model predicts that the evolution of worker polymorphism can be hampered if signalling of nutritional state by larvae is too costly, which prevents the evolution of larval signalling. This could be tested empirically by comparing the costs for larval signal production in species with polymorphic workers with the cost in species having monomorphic workers. Unfortunately, currently very little is known about the costliness of larval signals in social insects [39].

### (c) Task specialisation and division of labour

In our model, worker polymorphism evolves only under accelerating foraging returns of increasing body size, and workers evolve task specialisation by body size, with small workers nursing the brood and large workers foraging for resources (Fig. 2f). This is in line with division of labour models which typically predict that accelerating fitness returns of the level of specialisation are required for the evolution of division of labour [21–23], although such benefits of specialisation are sometimes apparently absent in empirical studies on task specialisation [61,62]. In our model, some degree of specialisation by body size also evolves under diminishing returns (Fig. 2c), although discrete worker polymorphism does not evolve. Similarly, some monomorphic ant species have been found to exhibit size-based task specialisation [63,64]. Body size might consequently function as a cue for dividing labour among the members of a colony in the absence of other cues, even if there are no accelerating efficiency benefits of task specialisation.

### (d) Larval signalling

The effect of larval signalling on worker division of labour and within-colony dynamics remains a little studied subject in social insect research [39]. In our model, worker polymorphism emerges due to the interplay of the evolved signalling strategies of larvae and the worker response to these signals. Larvae evolve to produce a nutrition-dependent signal and workers evolve to preferentially feed larvae with stronger signals. According with these model predictions, larval signalling has been demonstrated in a range of social insect species, as well as preferential feeding of signalling larvae by workers [41–45]. A crucial prediction of our model is that larval signalling strategies leading to worker polymorphism evolve to be non-monotonic with respect to larval condition. This prediction could be tested empirically by reconstructing the signalling reaction norms of larvae as a function of their nutritional state. If such non-monotonic signalling strategies are confirmed empirically, this would be evidence in favour of the model prediction that worker polymorphism emerges due to an interaction of larval signalling and worker response to the signals.

In our model, the fitness interests of all individuals within a colony are aligned – there is no conflict about sex allocation or male production. However, if such conflicts existed this could open up the opportunity for (allo)parent-offspring conflict, sibling-sibling conflict and dishonest signalling [53,54,65], which could affect body size distributions within colonies and might alter our model predictions. For instance, in species where workers can produce male eggs, larvae might evolve to signal more selfishly to improve their future reproductive abilities. In our model, polymorphic workers emerge because larvae that have obtained large amounts of food evolve to signal less than their siblings who have received less food. Selfish larvae that signal to obtain resources for producing male brood might not evolve such a strategy but signal at high levels even after obtaining a lot of food. Therefore, larval signalling strategies that lead to worker polymorphism might not evolve in such a scenario. In species with high levels of queen polyandry, policing of worker reproduction is more intense, leading to the evolution of worker sterility [66–68]. This in turn might lead to the evolution of less selfish larval signalling strategies and thus enable the evolution of worker polymorphism. Consequently, queen polyandry could have a facilitatory effect on the evolution of worker polymorphism, not by increasing body size diversity within a colony, but rather by leading to the evolution of worker sterility and less selfish larval signalling.

## 5. Conclusion

The interplay between larval signalling behaviour and worker response to the larval signals remains a poorly studied subject in social insect research [39]. We approached this issue theoretically by demonstrating in an evolutionary individual-based simulation model that the interplay between larval signalling and worker behaviour can lead to the evolution of mechanisms that result in the production of workers of distinct size classes and thus to worker polymorphism. This self-reinforcement mechanism resembles a nutrition-dependent developmental switch for caste determination, as ubiquitously found across eusocial insects. Our model therefore might provide a step towards a better understanding of social interactions of individuals of different developmental stages within social insect colonies, yielding some empirically testable predictions that might inspire future research.

## Data accessibility

Simulation model code, data analysis scripts and simulation data are available under https://doi.org/10.34894/EOHS8J. Temporary preview link for review: https://dataverse.nl/privateurl.xhtml?token=2e4136d8-7edc-4681-9d9f-24a98b540060.

## Author contributions

JJLO, IP and JJK conceptualized and implemented the model. JJLO and JJK analysed the model. JJLO, IP and JJK wrote the manuscript.

## Acknowledgements

We thank Thijs Janzen for helping us to speed up the simulations. We would like to thank the Center for Information Technology of the University of Groningen for providing access to the Peregrine high performance computing cluster. JJLO was supported by the EMJMD scholarship of the European Commission by the Erasmus Mundus Master Programme in Evolutionary Biology (MEME) and the DFG Emmy Noether Program #511474012. JJK was supported by an Adaptive Life grant by the University of Groningen.

