## Supplement for "Coevolution of larval signalling and worker response can trigger developmental caste determination in social insects"

### Modelling reaction norms with natural cubic spline functions

We model flexible smooth reaction norms with natural cubic spline functions. These functions consist of connected cubic polynomials but they are linear on both tail ends. We follow the definition of a natural (or restricted) cubic spline function by Harrell [1]. A natural cubic spline function has $k$ knots, which are the connection locations of the cubic polynomials with respect to $x$ (in the main manuscript, we set $k = 5$). The knot locations of the $k$ knots are $t_1, \ldots, t_k$. The natural cubic spline function, consisting of the basis functions $B_i$, is then given by

$$f(x) = \beta_0 B_0 + \beta_1 B_1 + \beta_2 B_2 + \cdots + \beta_{k-1} B_{k-1}, \quad (1)$$

where $B_0 = 1$ and $B_1 = x$. The remaining terms $B_2, \ldots, B_{k-1}$ are calculated by iterating over $j = 1, \ldots, k-2$ according to

$$B_{j+1} = (x - t_j)_+^3 - \frac{(x - t_{k-1})_+^3 (t_k - t_j)}{t_k - t_{k-1}} + \frac{(x - t_k)_+^3 (t_{k-1} - t_j)}{t_k - t_{k-1}}, \quad (2)$$

where $(\ldots)_+$ indicates that a term is set to 0 when it is evaluates to a number below 0. The natural cubic spline function can also be written in matrix notation.

$$f(x_i) = \begin{bmatrix} B_0(x_1) & B_1(x_1) & B_2(x_1) & \cdots & B_{k-1}(x_1) \\ B_0(x_2) & B_1(x_2) & B_2(x_2) & \cdots & B_{k-1}(x_2) \\ \vdots & \vdots & \vdots & \ddots & \vdots \\ B_0(x_n) & B_1(x_n) & B_2(x_n) & \cdots & B_{k-1}(x_n) \end{bmatrix}_i \begin{bmatrix} \beta_0 \\ \beta_1 \\ \beta_2 \\ \vdots \\ \beta_{k-1} \end{bmatrix} \quad (3)$$

Here, the matrix is a so-called basis matrix of the natural cubic spline function. The column vector $\beta$ contains the parameters $\beta_0, \ldots, \beta_{k-1}$ for the natural cubic spline function. The basis matrix subscript $i$ signifies that the $i$-th row of the basis matrix is used for the multiplication with the column vector $\beta$. In our simulations, we assume that the knot locations are evenly distributed within the range of $x$. The basis matrix is precalculated on equally-spaced intervals at locations $x_i$ at initialisation for a given resolution of the natural cubic spline function for a given range of $x$ (in the main manuscript, we set the resolution to 100). In the case of the reaction norm for larval signalling, we set the range of $x$ between $X_{\min} > 0$ and $X_{\max} = 5$; i.e., workers are larger than body size 0 and their maximum possible body size is 5. The natural cubic spline function parameters in column vector $\beta$ are the evolving gene values. In the reaction norm for larval signalling, we initialised all elements of $\beta$ with 0. Consequently, larvae initially produce no larval signals at all, regardless of their nutrition level $X$. (When larvae develop to workers, their nutrition level determines their body size $X$. Thus, the larval nutrition level is also $X$.) When evaluating the reaction norm for individual larva, we identify the two closest prespecified $x_i$ to the nutrition level $X$ of the larva and select one of these. Subsequently, we multiply the column vector $\beta$ with the $i$-th row of the basis matrix. This yields a single value, which is the phenotypic value (in this case, the signalling strength) for a particular value of $x$.

To obtain a multi-dimensional natural cubic spline surface, we construct two basis matrices. We then obtain a new basis matrix of $x^2$ rows and $k^2$ columns by taking the tensor product of each row of the first basis matrix with each row of the second basis matrix. The column vector of gene values now has a size of $\beta^2$. Again, the multiplication of the $i$-th row of the new basis matrix with the gene value column vector reduces to a single phenotypic value for a given value of $x$ and $y$. When evaluating the reaction norm for a worker, we again randomly select between the two precalculated values of $x$ and $y$, respectively, closest to the actual values of $x$ and $y$. Again, this yields the row index $i$. The values of $x$ again have the range of body sizes from $X_{\text{min}} > 0$ to $X_{\text{max}} = 5$. The values of $y$ reflect the sum of larval signals, which ranges from $S_{\text{min}} = 0$ to $S_{\text{max}} = 1000$. We again assume that the knot locations are evenly distributed within those ranges. We logistically transform the phenotypic value obtained from evaluating the natural cubic spline surface to a foraging probability. We initialise the genetic values with values of 0. After logistic transformation, this results in a foraging probability of 0.5, regardless of the larval signalling strength and the worker's body size.

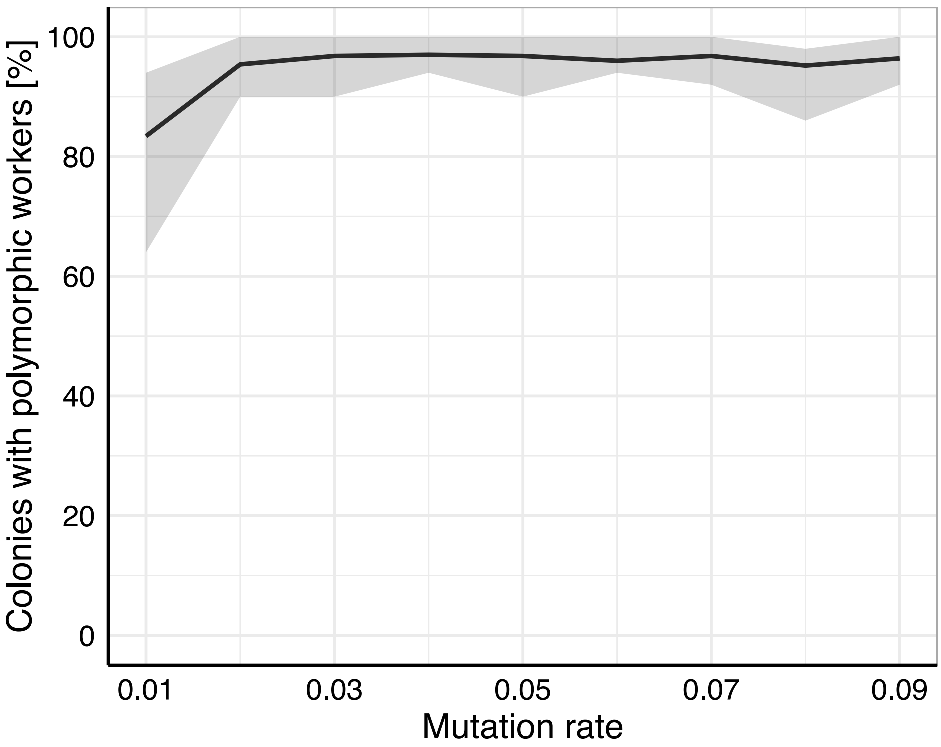


**Figure S1.** The effect of mutation rate on the evolution of worker polymorphism. When the mutation rate is low, worker polymorphism is slightly less prevalent, since a larger number of generations would be required to reach the same evolutionary equilibrium. We simulated a range of 0.01 to 0.09 in steps 0.01 of mutation rates ($m$), each with 10 replicate simulations. Black lines show the mean percentage of colonies with polymorphic workers and grey areas the range across replicates. We assume accelerating foraging returns of increasing body size ($b$ = 1.5).

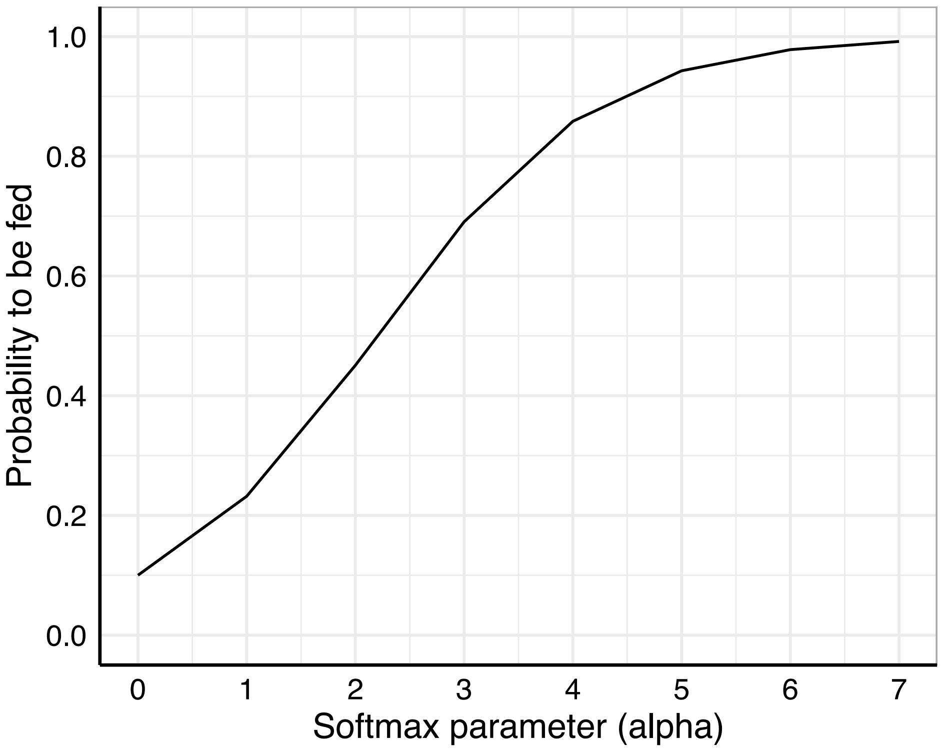


**Figure S2.** The effect of the softmax parameter ($\alpha$) on the probability of a larva to receive food. This parameter affects eq. 3 from the main manuscript: $P_l = \frac{\exp(S_l\alpha)}{\sum_{i=1}^{L}\exp(S_i\alpha)}$. We here assume a scenario with 10 larvae where one larvae produces a signal that is stronger by 1.0 than the signal produced by each of the other larvae. The y-axis shows the probability that the larva producing the stronger signal receives food.

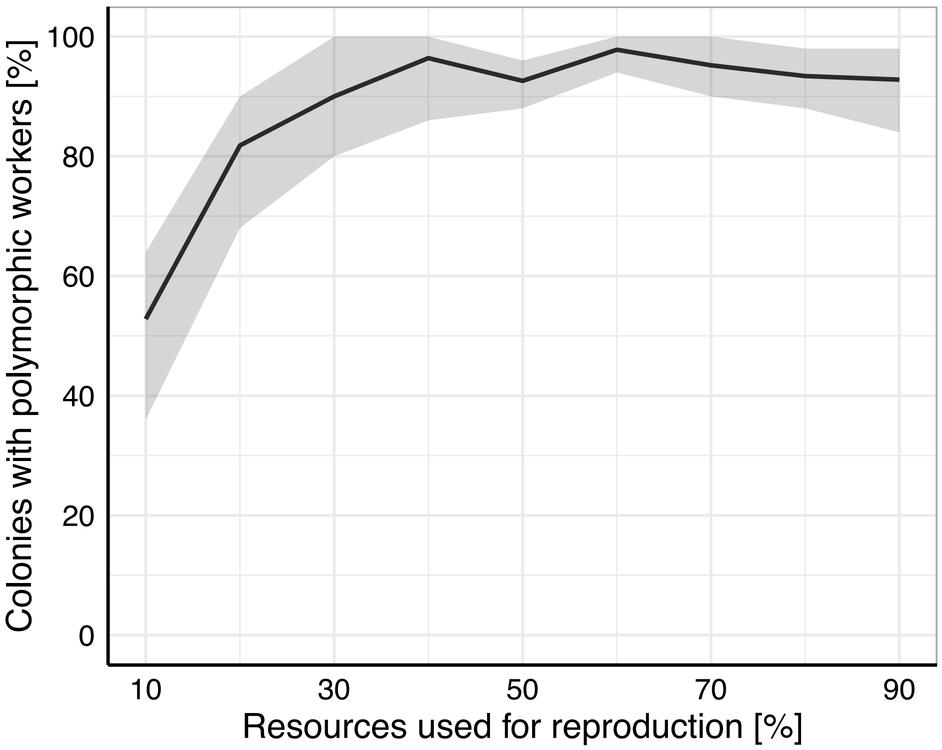


**Figure S3.** The effect of percentage of resources used for reproduction on the evolution of worker polymorphism. We simulated a range of 0.1 to 0.9 in steps 0.1 of the proportion of resources used for reproduction ($r$), each with 10 replicate simulations. Black lines show the mean percentage of colonies with polymorphic workers and grey areas the range across replicates. We assume accelerating foraging returns of increasing body size ($b = 1.5$).

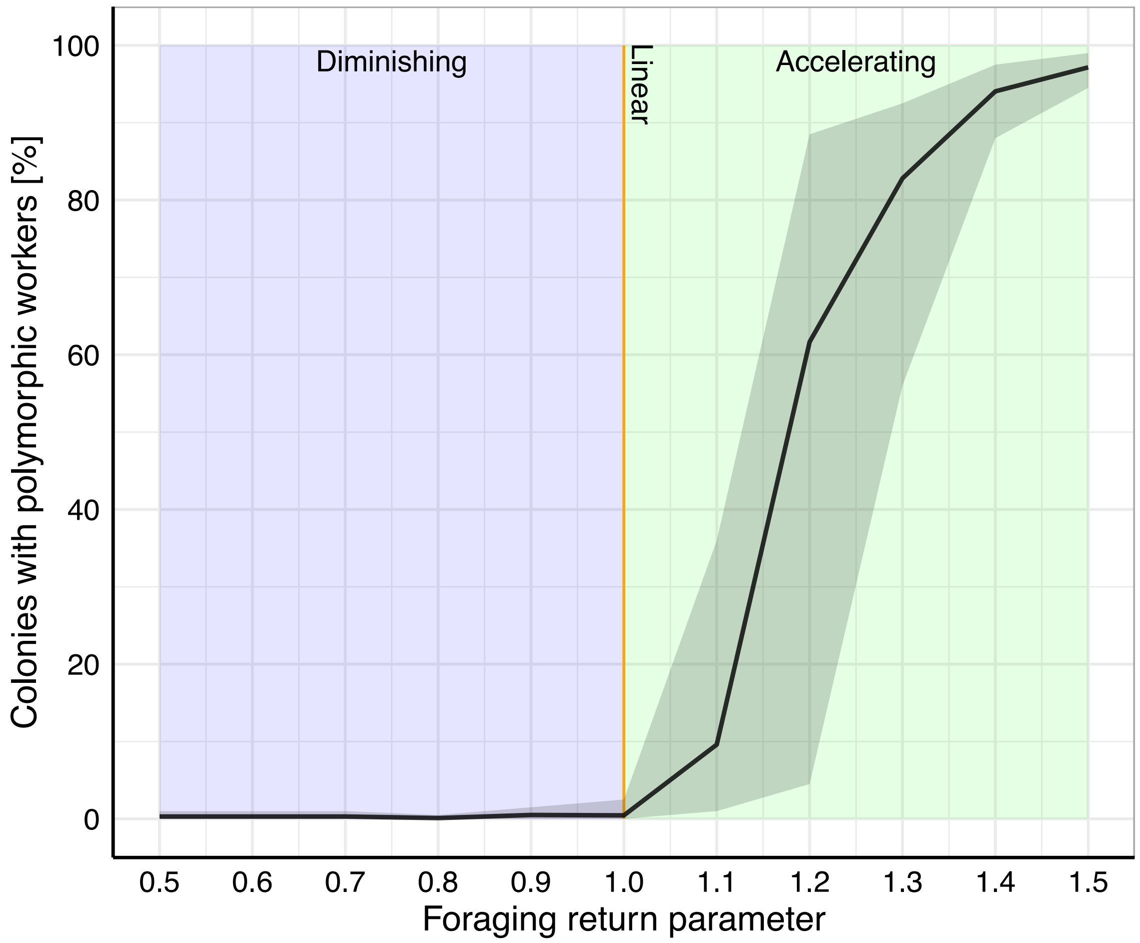


**Figure S4.** The effect of the foraging return parameter ($b$) on the evolution of worker polymorphism. This parameter affects the shape of eq. 2 from the main manuscript: $F = aX^b$. If the foraging return parameter is < 1 (purple area), then foraging returns diminish with increasing body size. If the foraging return parameter is exactly 1, then foraging returns increase linearly with increasing body size (orange line). If the foraging return parameter is > 1, then foraging returns accelerate with increasing body size. We simulated a range of 0.5 to 1.5 in steps of 0.1 of the foraging return parameter, each with 10 replicate simulations. The black line shows the mean percentage of colonies with polymorphic workers and the grey area the range across replicates.

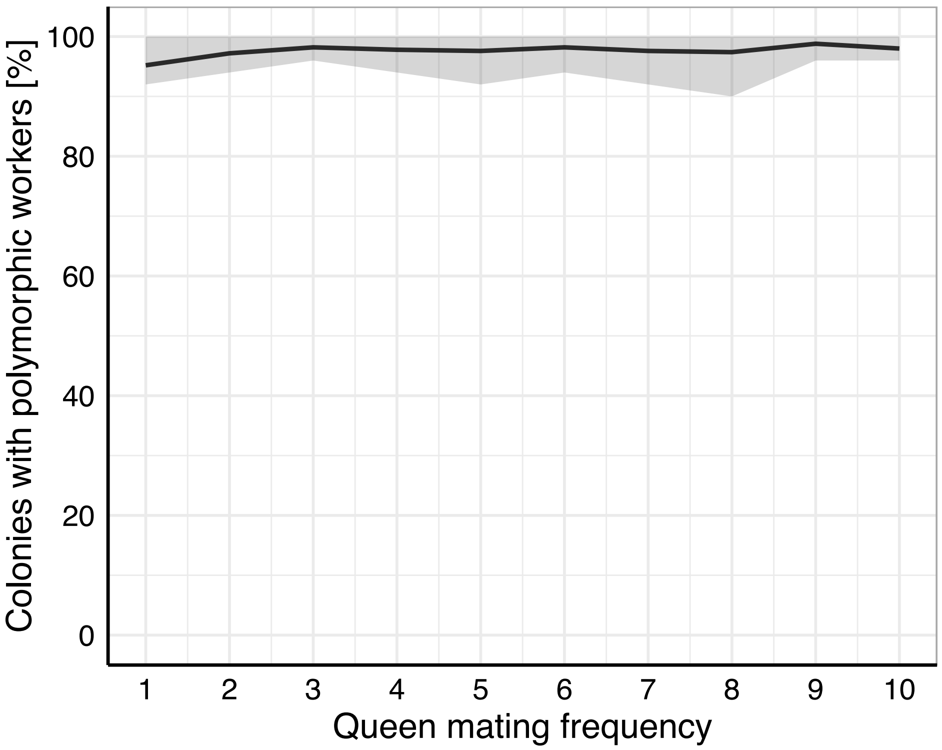


**Figure S5.** The effect of queen mating frequency on the evolution of worker polymorphism. We simulated a range of 1 to 10 in steps 1 of queen mating frequency ($n$), each with 10 replicate simulations. Black lines show the mean percentage of colonies with polymorphic workers and grey areas the range across replicates. We assume accelerating foraging returns of increasing body size ($b = 1.5$).

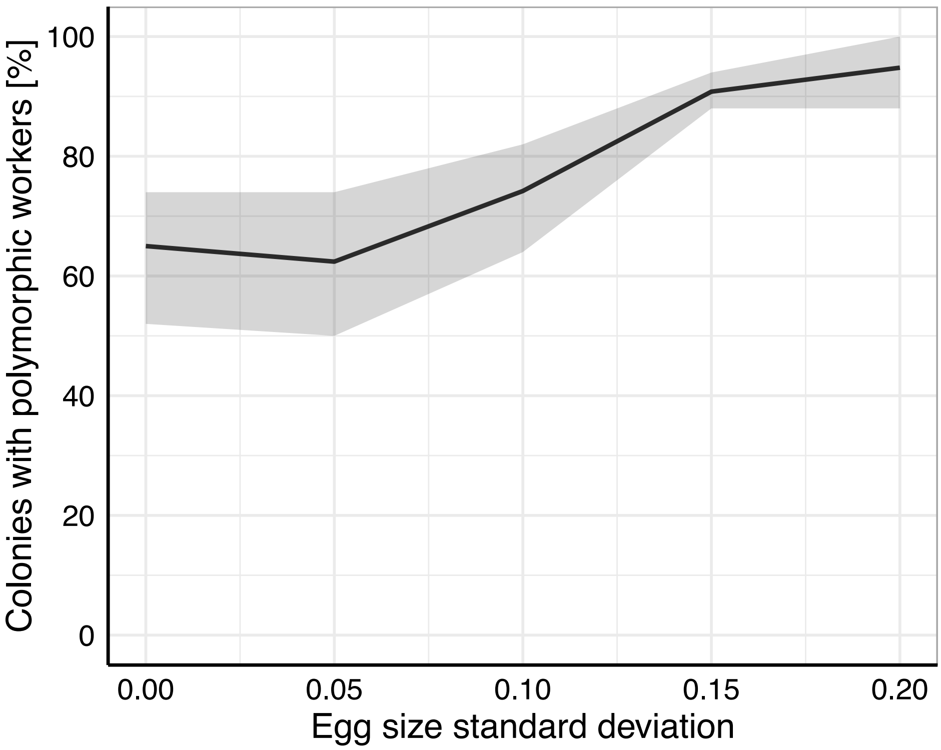


**Figure S6.** The effect of variation in egg size on the evolution of worker polymorphism. Worker polymorphism evolved in the absence of egg size variation. However, it is more prevalent when egg size variation occurs. We simulated a range of 0.00 to 0.20 in steps 0.05 of egg size standard deviation ($\sigma_{\text{egg}}$), each with 10 replicate simulations. Black lines show the mean percentage of colonies with polymorphic workers and grey areas the range across replicates. We assume accelerating foraging returns of increasing body size ($b = 1.5$).
